# Development of novel signal and spike velocity analysis tools in peripheral nerve cuffs

**DOI:** 10.1101/2024.11.20.624516

**Authors:** Jonas Klus, Alexander J. Boys, Ruben Ruiz-Mateos Serrano, George G. Malliaras, Alejandro Carnicer-Lombarte

**Author notes:** Author to whom any correspondence should be addressed.

## Abstract

**Objective:** Peripheral nerve neurotechnologies hold significant promise as avenues for new closed-loop clinical treatments. However, analysis tools for nerve recordings – a key component of closed-loop nerve technologies – remain underdeveloped compared to brain-focused methods. This study introduces and explores the performance of two novel nerve signal analysis techniques which rely on a defining feature of peripheral nerve signals: the reliable conduction velocity of signals transmitted by axons in nerves.

**Approach:** We test the capabilities of the introduced cross-correlation and spike delay velocity analysis techniques both *in silico* on synthetic nerve signals and on *in vivo* nerve signals acquired from freely-moving rats.

**Main results:** Our findings show that both techniques can be successfully employed to extract transmission direction and velocity information from nerve cuff recordings. Notably, cross-correlation analysis can be employed to detect neural signals of very low signal-to-noise ratio, otherwise undetectable by typical spike detection approaches.

**Significance:** Our findings provide new techniques to both enhance detection and extract new information in the form of velocity data from nerve recordings. As axon signal conduction direction and velocity is tightly linked to neural function, these techniques can support new research into peripheral nervous system function and new therapeutic approaches driven by neural interfaces.

## 1. Introduction

Implantable neurotechnologies, in particular those applied to the peripheral nervous system, are emerging as promising therapeutic modalities for a range of conditions. In clinical practice, conditions such as epilepsy and depression are now being treated through implantable nerve stimulators, which modulate nerve activity to alleviate symptoms [1]. Other nerve stimulation-driven treatments, such as those targeting immune responses, are also showing significant potential [2–4]. Similar to other neurotechnology fields, therapies that combine recording with stimulation in nerves in the form of closed-loop neurotechnologies are rapidly gaining attention as powerful treatment modalities [5–7].

The addition of nerve recording capabilities in closed-loop nerve neurotechnologies enables real-time monitoring of the state of the nerve, allowing for dynamic adjustment of the therapeutic stimulation. Although nerve recording is technically more challenging than stimulation, advances in materials and fabrication techniques for implantable devices now enable reliable nerve recording in both anesthetized [8–11] and freely-moving awake conditions [12]. Recent work has shown that activity recorded from peripheral nerves correlates with clinically relevant parameters, including blood glucose levels in diabetes research [13] and bladder state in spinal cord injury research [9]. These findings underscore the broad potential for closed-loop therapeutics powered by nerve recordings.

Despite this promise, neural analysis tools for peripheral nerve recordings remain underdeveloped compared to those used for brain recordings. Techniques such as spike sorting, commonly used in brain recordings, may be less effective for nerve signals, particularly when recorded through cuff electrodes placed around the nerve’s epineurium where large numbers of axon signals at low signal-to-noise ratio (SNR) are expected to contribute to the recordings. The anatomy of peripheral nerves, however, presents a unique opportunity for the application of velocity-based analysis approaches. Nerves are composed of bundles of axon fibres with diverse diameters and degrees of myelination, which affects the conduction velocity of action potentials and reflects the fibre’s functional role. By positioning two electrodes along the nerve’s length, the propagation of these signals could be recorded and employed to enable more robust and informative analyses of electrophysiological data from peripheral nerves.

In this work, we investigate the application of velocity-based analysis techniques in peripheral nerve recordings. We focus on two strategies: cross-correlation of signal and spike delay velocity analysis. First, we assess the validity and accuracy of these techniques on a synthetic dataset of recorded nerve signals. Building on these insights, we develop improved velocity-based analysis methods and apply them to nerve recordings from freely-moving awake rat models.

## 2. Method

### 2.1. Synthetic nerve signal simulation

Synthetic nerve signals were generated using an object-oriented *Python 3*.*12*.*4* framework, incorporating custom-built classes to simulate recordings at two electrode sites spaced 2 mm apart. Additional classes were designed to handle spike generation and noise addition. The simulation utilized *NumPy (v1*.*26*.*4)* for numerical computations, *Pandas (v2*.*2*.*2)* for data management, *SciPy (v1*.*13*.*1)* for signal processing tasks like filtering, and *Matplotlib (v3*.*8*.*4)* for visualization. Spike velocities were predefined at 1, 5, 10, 30, 60, and 120 m/s, serving as a simplified model of different nerve fibre types [14,15]. Velocities were assigned randomly to each spike. During simulation trials, approximately 60 spikes per second were produced. Spikes were modelled using exponential functions that approximate extracellularly recorded spike shapes [16,17]. Details of these functions are provided in the Supplementary Fig. 1. The peak amplitude of spikes was adjusted using a scaling factor, which preserved the spike shape while proportionally increasing or decreasing the overall amplitude of both the negative depolarization peak and the positive hyperpolarization peak. This approach enabled the modelling of spikes with high amplitudes exceeding 7 times the root mean square (RMS) of the background noise, as used in our spike velocity analysis, as well as very low amplitudes near 1 RMS of the background noise, as used in our cross-correlation analysis.

**Figure 1.**
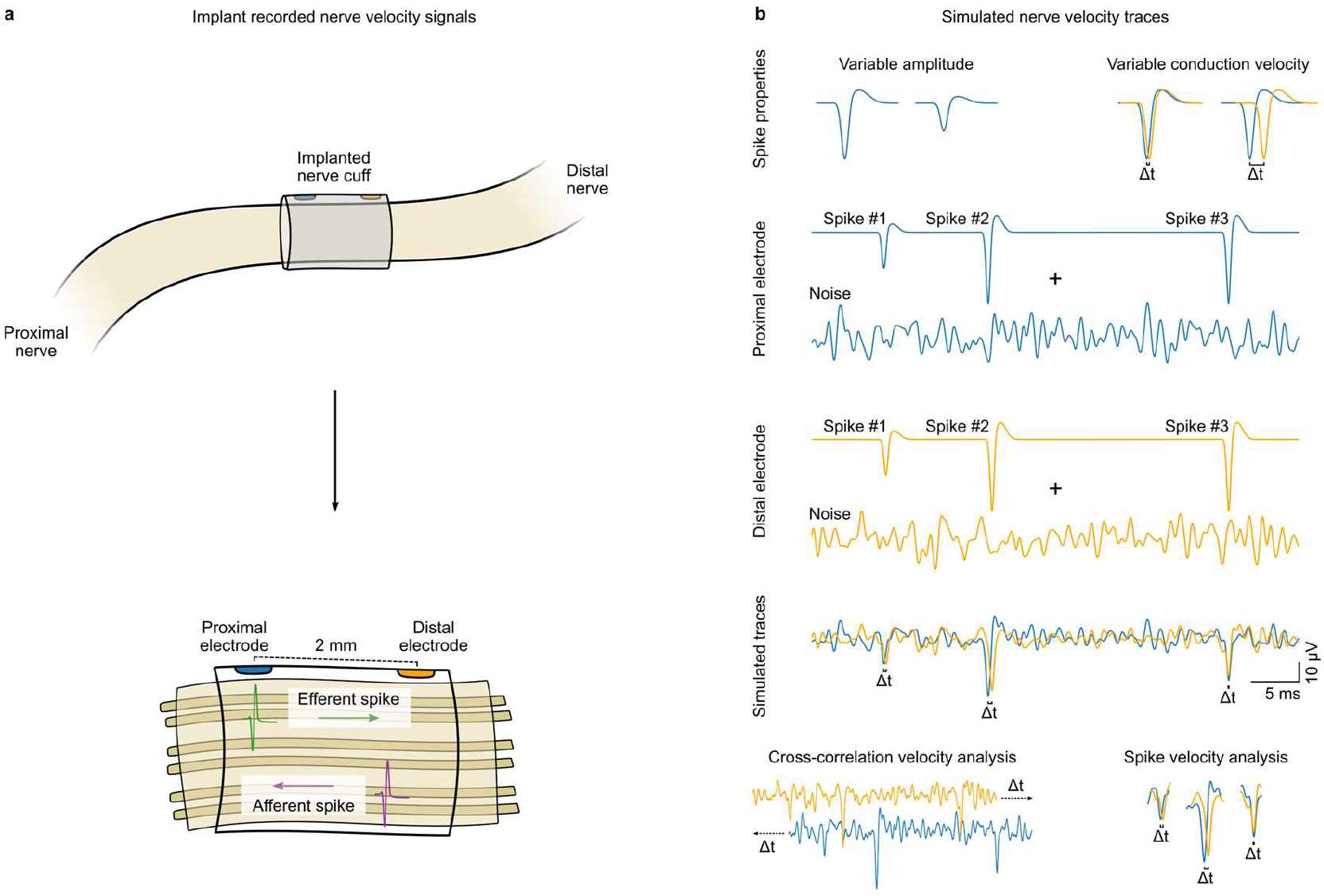
Simulation of nerve signals using a synthetic nerve cuff model. a) Schematic of the implanted nerve cuff electrode setup for recording action potential propagation in peripheral nerves. The cuff encompasses a section of the nerve with two electrode sites positioned 2 mm apart, allowing for the detection of efferent and afferent spikes, each representing distinct signal directions within the nerve bundle. b) Simulated nerve velocity traces demonstrating a range of spike properties. Top row: Representative spike waveforms illustrating different simulated spike amplitudes and conduction velocities. Middle rows: Simulated proximal (blue) and distal (yellow) electrode recordings, each composed of distinct spike events (Spike #1, Spike #2, Spike #3) combined with background noise to mimic real nerve recording conditions, and how these are superimposed showing spike alignment differences between the two recording sites (Δt), caused by the simulated time delay in recorded signals due to varying conduction velocities. Bottom row: two velocity analysis tools tested in this study: cross-correlation analysis of recorded trace segments, where the correlation between recordings is compared at different time delays; and analysis of spike within the trace, where the delay between spikes across recordings is compared.

The ground truth signal was initially created at 3 MHz to provide a high-resolution approximation of a continuous nerve signal, selected to facilitate straightforward calculations and minimize rounding errors during subsequent sampling to 30 kHz. This sampled rate was chosen as it represents the maximum digital sampling rate supported by common commercial electrophysiology hardware. Spikes were modelled to propagate across the 2 mm gap between electrodes (Fig. 1a), with the time delay between recordings calculated based on velocity and then converted into sample point differences across the two traces. The electrodes were assumed to be infinitesimally small points, simplifying the propagation model by focusing solely on the time delay between the two locations.

To simulate background noise in *in vivo* nerve recordings, we used *Python’s NumPy random*.*normal* function to generate white noise, drawing random samples from a normal (Gaussian) distribution. This noise was then filtered through a 5th-order Butterworth low-pass filter with a cutoff frequency of 1 kHz to reduce high-frequency components. The filtered noise was added to the spike traces, creating a more realistic total nerve signal (Fig. 1b).

A time array was generated for both the analogue (3 MHz) and the digital (30 kHz) signal, ensuring accurate spike and noise positioning over the trial. Data for each spike, including amplitude, velocity, and peak positions in both analogue and digital formats, were stored in a *Pandas DataFrame*, facilitating efficient tracking and analysis across trials. The final synthetic nerve signal combined spike traces with filtered noise (Fig. 1b), providing both high-resolution analogue and sampled digital versions. Each trial introduced unique spike and noise combinations, simulating real-world variability.

### 2.2. Signal cross-correlation

Cross-correlation analysis was conducted to evaluate the similarity between two signals at various time lags, aimed at detecting time delays between spikes recorded at the two electrodes spaced 2 mm apart. The cross-correlation of two signals, *f(t)* and *g(t)*, with *τ* as the time lag, is defined as

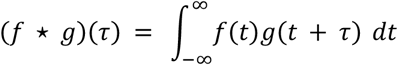

This calculation provides a measure of similarity between the two signals as one is shifted in time relative to the other. Using *NumPy’s correlate* function in *full* mode, the cross-correlation was computed for all possible time lags, from the maximum negative lag (where spikes in the second signal lead those in the first) to the maximum positive lag (where spikes in the first signal lead those in the second), capturing the entire range of time-lagged correlations.

For the whole signal cross-correlation analysis (Fig. 3), a time array was generated to span the full range of possible time lags, encompassing both negative and positive values. Analyses and graphs were presented with the x-axis scaled in milliseconds (Fig. 2, 3), aligning with the expected propagation times for nerve signals. This scaling enabled direct interpretation of time delays in terms of spike propagation velocity. To assess low-amplitude spike detection, cross-correlation analyses were conducted using traces ranging from 100 milliseconds to 30,000 milliseconds (Fig. 3p). For a single analysis segment, ten spikes were added at equidistant intervals to an array of 300,000 zero-valued sampling points (Fig. 3a-c), followed by the addition of random noise to create a realistic composite signal (Fig. 3e-g). For longer traces, multiple spike-only segments were concatenated, and random noise was applied across the entire combined signal (Fig. 3m-p). This approach enabled the examination of cross-correlation performance in detecting low-amplitude spikes over varying signal lengths.

**Figure 2.**
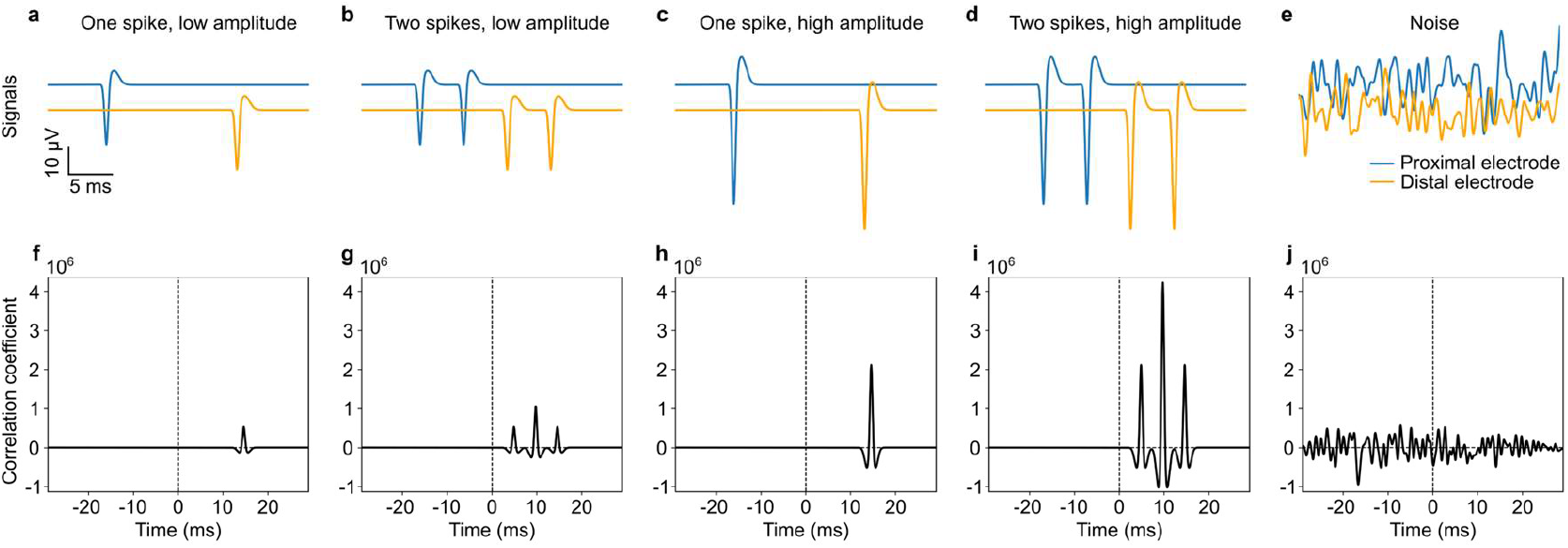
Effect of spike properties on cross-correlation analysis. (a–d) Simulated proximal (blue) and distal (yellow) electrode traces illustrating nerve signals with different spike characteristics: single low-amplitude spike (a), two low-amplitude spikes (b), single high-amplitude spike (c), and two high-amplitude spikes (d). (e) Noise-only signal. (f–j) Corresponding cross-correlation plots for each condition (a–e), showing correlation over time delay. In signals with multiple spikes, the additive effect of spikes increases the cross-correlation magnitude (b, g). Higher amplitude spikes create more pronounced peaks due to their multiplicative effect (c, h). Combined effects of multiple and high-amplitude spikes produce both additive and multiplicative impacts on cross-correlation (d, i). Noise produces no distinct peaks, with correlation values fluctuating around the mean (e, j). Correlation coefficient zero has been offset for easier comparison across graphs.

**Figure 3.**
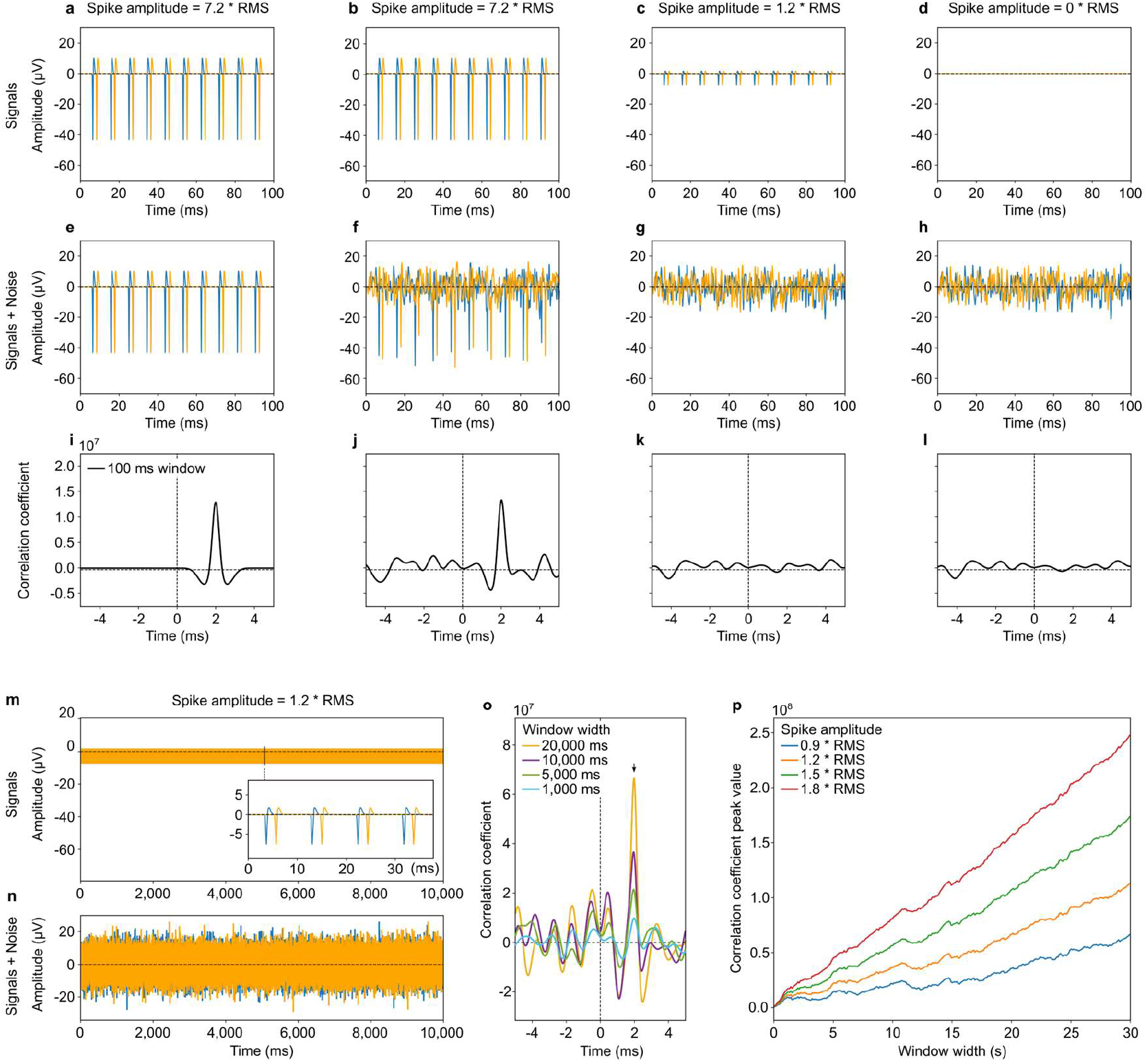
Cross-correlation analysis of signal detection under varying spike amplitude and signal window lengths in noisy environments. (a–d) Simulated nerve signals with different spike amplitudes: high-amplitude spikes at 7.2*RMS of noise (a, b), low-amplitude spikes at 1.2*RMS of noise (c), and no spikes as a control (d). (e–h) Addition of noise to each condition in (a–d), with the noisy signals used to evaluate the effect of noise on spike detection. No noise added in (e) as a control. (i–l) Cross-correlation plots for each signal in (e–h) using a 100 ms window. High-amplitude spikes (7.2*RMS) produce a distinct peak in the cross-correlation, even in noisy conditions (j), whereas low-amplitude spikes (1.2*RMS) are obscured by noise, resulting in a flat cross-correlation pattern similar to the noise-only condition. (m–o) Analysis of detectability over extended time windows. (m) Simulated signal with spike amplitude of 1.2*RMS of noise over 10 s. Inset highlights spikes within a shorter interval for clarity. (n) Corresponding signal with noise added. (o) Cross-correlation results over a range of window widths, showing that increasing the window length from 1,000 ms to 20,000 ms allows produces a higher correlation peak (black arrow). (p) Plot of correlation peak values as a function of window length for spikes with different amplitudes. Higher-amplitude spikes require shorter windows to yield higher amplitude correlation peaks.

### 2.3. Peak search and velocity estimation

Two spike velocity estimation methods—peak interval velocity and cross-correlation coefficient—were employed to assess nerve signal velocity across different RMS thresholds. For both methods, peaks were first identified using a hill climbing algorithm. Only negative spikes were considered, as we simulated extracellular recordings [16]. The peak velocity method then calculates velocity based on the difference in sampling points between peaks. The cross-correlation coefficient method computes time-shifted cross-correlation coefficients around the identified peaks to determine velocity.

The RMS threshold in this context is used to detect peaks in the signal that represent potential spikes. Calculated as a multiple of the signal’s RMS, this threshold sets a minimum amplitude that a data point must exceed to be classified as a spike [18]. By defining the threshold as an RMS multiple, this approach dynamically adapts to signal variability, distinguishing spikes from background noise. This method assumes that spikes are sporadic, meaning their influence on the RMS calculation is minimal and the RMS primarily reflects the noise floor. Higher thresholds reduce the likelihood of false positives by requiring larger deviations from the mean, while lower thresholds increase sensitivity to smaller amplitude spikes, though with a higher risk of false positives. The optimal threshold depends on the specific study objectives. Thus, different thresholds were employed in our analyses to test changes in performance when accommodating varying detection needs.

The peak detection uses a hill climbing algorithm to identify corresponding spikes in two signals recorded at electrode sites spaced 2 mm apart (Fig. 1a). Hill climbing works by comparing each data point to its immediate neighbours to locate local maxima, moving point by point until a peak is identified. In the initial signal, peaks are detected using a threshold set as a multiple of the signal’s RMS. Once a peak is detected, a search window of 100 samples is established around the peak to locate a corresponding spike in the second signal for the sampled signal. For interpolated signals, this search window is scaled by the upsampling factor, which in our simulations is 100 as described below, resulting in an effective search window of 100 * 100 = 10,000 samples.

The peak velocity is then calculated by determining the sampling point delay between the detected peaks in the two signals. The velocity calculation follows

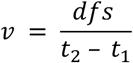

where *d* is the electrode distance (2 mm), *f* is the sampling rate (30 kHz), *s* is the search scaler (1 for simple inter-spike interval and 100 for sinc and cubic spline interpolation as detailed below), and *t*_*1*_ and *t*_*2*_ are the detected peak times in the first and second signals, respectively. The calculated velocity is compared to the true velocity and classified as correct if it falls within ±40% of the true speed, which implies that the direction is correctly identified as well. This 40% margin was arbitrarily selected as it allows for approximate classification of spikes to specific fibre types, preserving clear separation between modelled true velocities, even in the presence of modest velocity estimation errors. However, we also created plots displaying in more detail the magnitude of any errors in velocity estimation.

The cross-correlation coefficient method provides an alternative velocity estimation by analysing the similarity between two signals as a function of time lag. After initial peak detection using hill climbing, a 100-sample window is extracted around each peak in both signals. The cross-correlation is calculated using *NumPy’s correlate* function, with the time lag at the maximum correlation value giving the delay *Δt* between spikes. The velocity for the corresponding spikes is then calculated using

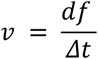

where *d* is the electrode spacing (2 mm), *f* is the sampling rate (30 kHz), and *Δt* is the sample delay. The velocity is again classified as correct if it falls within 40% of the true speed. In contrast to its earlier application (Fig. 2-3), cross-correlation here is employed to estimate delays between individual spike pairs, rather than whole signal traces.

### 2.4. Signal interpolation

In addition to testing two velocity estimation methods, the effect of interpolation to increase sampling rate on spike velocity estimation was also tested. Specifically, two interpolation techniques—sinc interpolation and cubic spline interpolation—were used to enhance signal resolution before performing peak search and peak interval velocity estimation.

The sinc interpolation method, which refines time resolution through upsampling and reconstructs the continuous signal, is defined as

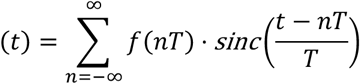

where *f(nT)* are discrete sampling points, *T* is the sampling interval, and *sinc(x)* is *sin(πx) / (πx)* [19]. The nerve signal is converted to the frequency domain via Fast Fourier Transform (FFT), zero-padded for higher resolution, and transformed back to the time domain with 100x finer detail.

The cubic spline interpolation method improves time resolution by fitting a piecewise cubic polynomial through the data points, allowing for smooth signal reconstruction. It constructs a continuous signal by interpolating between sampled points using cubic polynomials. For each interval [x_i_, x_i+1_], the cubic spline is defined as

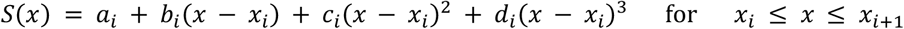

where x_i_ and x_i+1_ are consecutive discrete sampling points, a_i_, b_i_, c_i_, and d_i_ are coefficients determined from the data points, and x is any point within the interval [x_i_, x_i+1_]. This method generates a smooth curve between each pair of points, preserving continuity in both the first and second derivatives [20]. Like sinc interpolation, we used cubic spline interpolation to enhance signal resolution by a factor of 100x, enabling more accurate spike detection.

### 2.5. Spike velocity estimation performance evaluation metrics

Performance evaluation for all methods involved simulations across RMS thresholds from 2.0 to 7.0 in 0.05 increments, with 100 runs per threshold to ensure statistical robustness. Several metrics were calculated as percentages of all ground truth spikes:

- *A – True Spike, True Velocity:* Percentage of correctly detected spike pairs with correct velocity (within ±40% of the true speed).
- *B – True Spike, False Velocity:* Percentage of correctly detected spike pairs but with incorrect velocity.
- *C – Missed Spike:* Percentage of true spike pairs not detected.
- *D – False Spike:* Percentage of detected spike pairs without true corresponding spikes.

This categorization meant that A, B, and C groups summed to 100% of ground truth spikes. The error group *B – True Spike, False Velocity* can be further subdivided into spikes with the correct direction but incorrect velocity, spikes with incorrect direction and velocity, and spikes with no sampling point difference, causing the velocity to be calculated as infinite (Fig. 5).

### 2.6. *In vivo* validation

All *in vivo* procedures were done in accordance with the UK Animals (Scientific Procedures) Act, 1986. Work was approved by the Animal Welfare and Ethical Review Body of the University of Cambridge and was approved by the UK Home Office (project license number PP5478947). Experiments were conducted on Lewis rats (∼150g, ∼9 weeks of age) (Charles River, UK). Rats were group-housed in individually ventilated cages with ad libitum access to food and water for the duration of the study.

Surgical implantation of devices was carried out under isoflurane anaesthesia (2.25% v/v in medical oxygen). Body temperature was monitored and maintained using a thermal blanket. An ultraconformable parylene-C device containing 100 by 100 μm PEDOT:PSS microelectrodes divided into three cuffs was implanted into the right forearm of rats and around the median, ulnar, and radial nerves. The microelectrodes were divided into distal and proximal rings of three or four microelectrodes which were used to perform the velocity analyses in this work. The devices, together with a ground cable connected to screw drilled into the space above the cerebellum, were externalized via a head-mounted access port. Analgesia was provided for two days following surgery (Metacam, oral suspension), and were kept under post-operative observation for three days following surgery. The custom-fabricated devices and implantation locations used were the same as those previously used [12].

Electrophysiology recordings were carried out in awake rats three to five days post-implantation. Implanted devices were connected to an acquisition system (RHS Stim/Recording System, Intan Technologies). Nerve recordings were performed while animals were placed on a platform on which they were free to move, at 30 kHz sampling rate. Once acquired, these recordings were bandpass filtered (300 to 2000 Hz), and referencing was performed. For referencing, the signal of either a proximal or distal microelectrode had the average signal of its corresponding proximal or distal ring of microelectrodes subtracted. This removed correlated noise while preserving any delays in signal seen between proximal and distal electrodes.

Cross-correlation coefficient calculation was performed similarly to that used with synthetic data. In addition to this, a method was developed to visualize cross-correlation coefficient evolution over time in recordings in the form of heatmap plots (e.g. Fig 6b, middle). These plots were generated by performing cross-correlation calculations at different intervals along the recording. Intervals of 50 ms (Fig. 6b-c) or 0.5 ms (Fig. 6d) step sizes were used to create the plots shown. The averaging window (time window over which cross-correlation was calculated at each step) was varied as indicated in the heatmap plots. Spike velocity calculation was performed in the same manner as used with synthetic data, with measured spike delays between -2 and +2 ms grouped into histograms with bin size 0.1 ms (Fig. 7). Within the central bin, spikes with a delay of exactly 0 ms were highlighted in red.

## 3. Results

### 3.1. Synthetic nerve signal simulation

Nerves are composed of bundles of axons, each axon being capable of conducting action potential signals at a particular velocity. These action potentials involve changes in membrane potential which can be recorded extracellularly. While action potential spike amplitudes may be large when recorded in close proximity to neurons or axons, recorded amplitude decreases as the distance between the membrane and recording electrodes increases [21,22]. Recordings with cuff electrodes generally capture action potentials with amplitudes on the order of microvolts, alongside baseline electrical noise. Using the characteristics of nerve cuff recordings – frequent spikes with low SNR and varying conduction velocity (Fig. 1a) – we designed a synthetic model to approximate peripheral nerve cuff recordings (Fig. 1b).

The synthetic nerve recording model was designed with simplified spike shapes and noise patterns to highlight key patterns observed in real data, enhancing clarity and interpretability. This approach was not intended to capture the full complexity of real nerve signals but instead focused on creating a tool that allows for straightforward analysis of the results. To simulate above- and below-threshold spikes, two types of spikes were generated with different amplitudes (Fig. 1b). Each spike was modelled with three simple exponential functions, selected for their similarity to extracellular spike shapes, then normalized and scaled by RMS values to achieve realistic approximations (Supplementary Fig. 1). By limiting the model to two distinct spike shapes, we aimed to strike a balance between biological realism and model simplicity, which aids in the interpretation of the results.

The spatial setup of the model was designed with a 2 mm distance between the two recording electrodes (Fig. 1a). This separation reflects a realistic configuration for a single multielectrode cuff implant [9,12,23,24], and aligns with the electrode configuration used in our *in vivo* model validation. Using a single cuff also reduces complications related to imprecision from animal movement or accurate implant location, which can occur in dual-cuff setups.

The digital sampling rate in our model was set at 30 kHz, which represents the maximum rate supported by common commercial electrophysiology hardware This sampling rate also matches the one used in our in *vivo* model validation. The analogue signal was modelled at 3 MHz, providing 100 ground truth sampling points between each digital sampling point, thereby closely approximating a continuous signal for our experimental purposes. Simulated spike velocities were set at 1, 5, 10, 30, 60, and 120 m/s, serving as a simplified model of different nerve fibre types [14], including slower-conducting C fibres and Aδ fibres associated with pain and temperature sensory transmission, as well as faster-conducting Aα and Aβ fibres involved in motor as well as sensory proprio- and mechano-receptive functions [15]. This discrete set of velocities ensures an unambiguous ground truth for analysis. Of particular importance, 60 m/s represents the highest velocity detectable with the 30 kHz sampling rate and 2 mm electrode spacing, corresponding to exactly one sample point difference. The 120 m/s velocity was included to simulate a detected spike ‘infinite’ velocity scenario, where no sample point difference would be detected in the discretely sampled signal.

To simulate realistic i*n vivo* conditions, white noise was generated and added to the spike signal. This white noise mimics the broadband noise frequently observed in experimental nerve recordings. A low-pass filter was applied to the white noise, ensuring that only relevant frequency components were preserved (Fig. 1b). High- and low-amplitude spikes were randomly generated and distributed across the trace to ensure a realistic yet simplified signal composition (Fig 1b). The random generation of spikes ensured variability consistent with *in vivo* recordings, avoiding systematic biases in the simulated signals.

### 3.2. Cross-correlation velocity analysis

Cross-correlation is a technique used to measure the similarity between two signals as one signal is shifter (positively or negatively) relative to the other. It is widely applied in various areas of signal processing and pattern recognition [25]. When a propagating signal conducted by a nerve such as a spike is recorded at two different locations by for example two electrodes, the same signal is captured with a time delay between two electrode recordings (Fig. 1b). By applying a cross-correlation analysis to the recordings of the two electrodes as a function of time delay, we can detect the co-occurrence of the spike, as well as the time lag at which it occurs. The position of the resulting peak in the cross-correlation function corresponds to the time shift with which the signal is recorded across the electrodes, which allows us to estimate the velocity of the propagating signal. While the effect of this analysis can be best pictured with the propagation of an individual neural spike, its application does not require individual spike identification or isolation within a recording for its application, eliminating the need for spike detection techniques which may be poorly suited to low SNR signals recorded from nerve cuffs.

Our simulations showed that cross-correlation provides a robust way to detect nerve signals and their velocity (Fig. 2). Compared to single spikes (Fig. 2a, f), multiple spikes of the same amplitude present in the signal have an additive effect on the cross-correlation function, increasing the overall correlation signal (Fig. 2b, g). In contrast, increasing the amplitude of individual spikes has a multiplicative effect on the cross-correlation, producing even more pronounced peaks (Fig. 2c, h). Both effects can occur simultaneously, resulting in combined additive and multiplicative influences on the cross-correlation (Fig. 2d, i). Random noise, unlike structured signals, does not follow a consistent pattern, and thus its cross-correlation fluctuates around the mean without producing any distinct peaks or identifiable structure (Fig. 2e, j). This contrast highlights the effectiveness of cross-correlation in distinguishing spikes from noise, ultimately improving the performance of velocity estimation in noisy neural signals. Another observation of note made from these simulations is that peaks in correlation coefficient produced by spikes were flanked by dips in coefficient (Fig. 2f-i). This occurs due to the biphasic nature of the spikes, where the negative depolarization phase of the spike coincides with the positive hyperpolarization, resulting in a negative correlation.

Having confirmed the ability of cross-correlation analysis to identify spikes among noise, we tested whether realistic nerve signals could be processed with this technique. We simulated trains of low or high amplitude spikes (Fig. 3a-d) combined with background noise (Fig. 3e-h) and analysed their resulting cross-correlation function (Fig. 3i-l). A train of high-amplitude spikes (7.2*RMS of noise) combined with a noisy signal over a short 100 ms trace, produced a clearly identifiable cross-correlation peak (Fig. 3j). In contrast, a train of low-amplitude spikes (1.2*RMS) which visually cannot be identified from among noise (Fig. 3g), produces no identifiable peak in its cross-correlation (Fig. 3 k), with the cross-correlation pattern resembling that of a noise signal without any spikes (Fig. 3l).

Given the additive effect of increasing spike counts on cross-correlation strength, we hypothesized that extending the recording window over which the cross-correlation was calculated could uncover low-amplitude spikes otherwise masked by noise. To test this, we gradually extended the window length from 100 ms to 30 s. As shown in Fig. 3m-o, the ground truth spike with a +2 ms delay, initially obscured by noise in the 100 ms window (Fig. 3k), produced a clearly distinguishable cross-correlation peak with a 10 s window and was clearly visible at 20 s (Fig. 3o). We examined how this effect generalised to various spike amplitudes and window widths (Fig. 3p). The correlation coefficient peak at +2 ms cross-correlation steadily increased with increasing window width, with the effect being more pronounced for higher amplitude spikes. Our results indicate that cross-correlation analysis can identify signals made up of low amplitude spikes. By averaging the signal over a longer time window, the random noise averages out, while the consistent patterns, such as spikes, become more prominent. This results in a progressive increase in the signal-to-noise ratio which can allow for the detection of even sub-threshold spikes.

### 3.3. Spike velocity analysis

Individual neural spike detection is commonly conducted in both brain and peripheral nerve studies to identify and classify spikes based on shape and amplitude. Researchers often focus on detecting spike occurrences and clustering them according to waveform characteristics [26–29]. Although our analysis strategy operates within the broader field of neural spike detection, our focus is different: we analysed time delays between spike pairs recorded at two electrodes in the peripheral nervous system to both more reliably identify neural spikes as well as gain insights into signal transmission velocities (Fig. 1b).

Using the synthetic nerve signal simulation, we studied how different processing approaches affect spike and velocity detection. More specifically, we explored the effect of different spike velocity estimation methods and signal interpolation techniques to enhance signal resolution. The first approach tested the identification of spike peaks using a hill climbing algorithm and estimation of velocity based on the sampling point difference between signals recorded at two sites within a nerve cuff, positioned 2 mm apart (Fig. 1a). We refer to this approach as ‘Sampled signal + Peak velocity’. This spike velocity estimation method was applied to a digitally sampled signal, simulating *in vivo* nerve recording acquisition. However, the small delay expected in spike pairs for the highest conduction velocities suggested that spike peaks could be misidentified, leading to incorrect velocity estimations (Fig. 4a-b). We therefore tested whether signal interpolation could be used as a way to increase velocity estimation accuracy (Fig. 4c). We tested two commonly used interpolation methods: sinc and cubic spline interpolation. After applying these, we again identified peaks using a hill climbing algorithm and estimated velocity based on sampling point difference between signals. We refer to these two methods as ‘Sinc interpolation + Peak velocity’ and ‘Spline interpolation + Peak velocity’. Finally, we tested whether a different method of estimating spike velocity performed differently. After identifying peaks using a hill climbing algorithm (applied over the signal with no interpolation) we estimated their velocity based on their cross-correlation. We refer to this method as ‘Sampled signal + CrossCoeff velocity’. It should be noted that this implementation of cross-correlation for velocity estimation is different from that of our earlier application (Fig. 2-3). In contrast to this earlier application, cross-correlation here is employed to estimate delays between individual spike pairs, rather than whole signal traces. Spike detection was employed over signal segments exceeding a given amplitude threshold crossing. These four tested methods provided a way to establish whether changes in spike processing approaches could have a noticeable impact on peak detection and velocity estimation without exhaustively exploring every possible approach.

To evaluate the performance of each method, we conducted simulations across a range of signal RMS thresholds for spike detection and various spike velocities representing different nerve fibre types [14,15] (1, 5, 10, 30, 60, 120 m/s), at an approximate spike rate of 60 spikes/s (Supplementary Fig. 2). Each spike had an equal probability of being either a low-amplitude spike (3.6*RMS of baseline noise) or a high-amplitude spike (just above 7.2*RMS of baseline noise). These properties are similar to spike-rich nerve recordings acquired by us in our later *in vivo* experiments, and were chosen so half of the spikes had amplitudes above the tested spike detection thresholds while the other half had amplitudes within this tested range (prior to the addition of noise). For every detected spike, we evaluated the performance of the various velocity estimation approaches dividing them into groups expressed as percentages of all ground truth spikes. These consisted of: A) ‘True Spike, True Velocity’: correctly detected spikes with correct velocity estimates (within ±40% of the true speed), B) ‘True Spike, False Velocity’: correctly detected spikes but with incorrect velocity estimates, C) ‘Missed Spike’: true spikes that were not detected, and D) ‘False Spike’: detected spike pairs that did not correspond to true spikes. This categorisation meant that A, B, and C groups summed to 100% of ground truth spikes. The category of correctly detected spikes with incorrect velocity estimates was further subdivided into spikes with the correct direction but incorrect velocity, spikes with incorrect direction and velocity, and spikes where no sampling point difference was detected, resulting in an infinite velocity calculation.

All four methods exhibited similar patterns of performance when tested on signals of all combined velocities (1 to 120 m/s) (Fig. 4d-g). As spike detection threshold increased, the fraction of correctly detected spikes (groups A and B) decreased, while missed spikes increased (group C). Notably, no false spikes (group D) were detected by any method except for a small fraction at the lowest thresholds ∼2*RMS, indicating that co-detection of signals across two electrodes represents a robust way of eliminating this undesirable component of spike detection. In terms of velocity estimation, while all methods generally produced a higher portion of correct velocity estimations (group A) than incorrect estimations (group B), ‘Sampled signal + CrossCoeff velocity’ performed the worst of all four methods (Fig. 4d-g). This was particularly evident when focusing on 4*RMS spike detection threshold (Fig. 4h, group A: 32.9%, 34.8%, 34.8%, 28.7%; group B: 15.6%, 13.7%, 13.7%, 19.7%, ‘Sampled signal + Peak velocity’, ‘Sinc interpolation + Peak velocity’, ‘Spline interpolation + Peak velocity’, ‘Sampled signal + CrossCoeff velocity’, respectively; group C: 51.5% for all, group D: 0.0% for all). ‘Sampled signal + Peak velocity’ seemed to underperform compared to the interpolation-using methods, but a more detailed analysis of the velocity error magnitudes (Fig. 4i-l) showed that this group produced the highest fraction of high accuracy velocity estimations (within -5% and +5% of real velocity) (Fig. 4i).

The detailed velocity error analysis also yielded other insights. Due to the 2 mm distance between proximal and distal electrodes in our simulation, some portion of high velocity spikes were expected to yield peaks coinciding exactly in time across proximal and distal electrode recordings. In particular, a 2 mm gap implemented on signals sampled at 30 kHz would make a single sample point peak delay be equivalent to a velocity of 60 m/s. Higher velocities such as 120 m/s (included in our simulations and representative of the fastest axon fibres) could lead to coinciding peaks which would be calculated as having infinite velocity. Consistent with this, tested methods applied over these 30 kHz sampled signals yielded a portion of infinite velocity spikes (Fig. 4i, l). Both methods utilizing interpolation vastly suppressed this type of error (Fig. 4j-k).

Having examined overall spike velocity estimation method performance, we sought to focus on whether these differences were particularly important for specific spike velocities. All methods performed best for the lowest velocity spikes (1 m/s), and worst for the highest velocity spikes (120 m/s), with performance overall decreasing with spike velocity (Fig. 5 and Supplementary Fig. 3-6). Similar to what was observed earlier, ‘Sampled signal + CrossCoeff velocity’ performed worse for all individual tested velocities (Fig. 5d and Supplementary Fig. 6). In contrast, differences in performance between the other three groups appeared to vary based on spike velocity. ‘Sampled signal + Peak velocity’ yielded a higher portion of high accuracy spikes (−5% to +5% of real velocity), particularly in the moderate velocity ranges (30 and 60 m/s) (Fig. 5a and Supplementary Fig. 3), although it also yielded the highest portion of very high error results (darker purple bars in Fig. 5) including spikes classified as infinite velocity. ‘Sinc interpolation + Peak velocity’ (Fig. 5b and Supplementary Fig. 4) and ‘Spline interpolation + Peak velocity’ (Fig. 5c and Supplementary Fig. 5) in turn performed better at the highest velocities (120 m/s) and classified no spikes as infinite velocity. Overall, both interpolation methods yielded practically identical results in both tests on all combined velocity spikes (Fig. 4) and specific velocity spikes (Fig. 5). Ultimately, our results showed that while all tested methods represent a viable approach to spike detection and velocity estimation, the choice should align with the study’s objectives. If detecting high velocities beyond 60 m/s is crucial, interpolation is the only viable approach, despite reducing the overall proportion of spikes identified with very low error. In research scenarios where both directionality accuracy and very low error rates are to be prioritised, and velocities beyond 60 m/s are not needed, the simpler sampled signal approach may be preferable.

**Figure 1.**
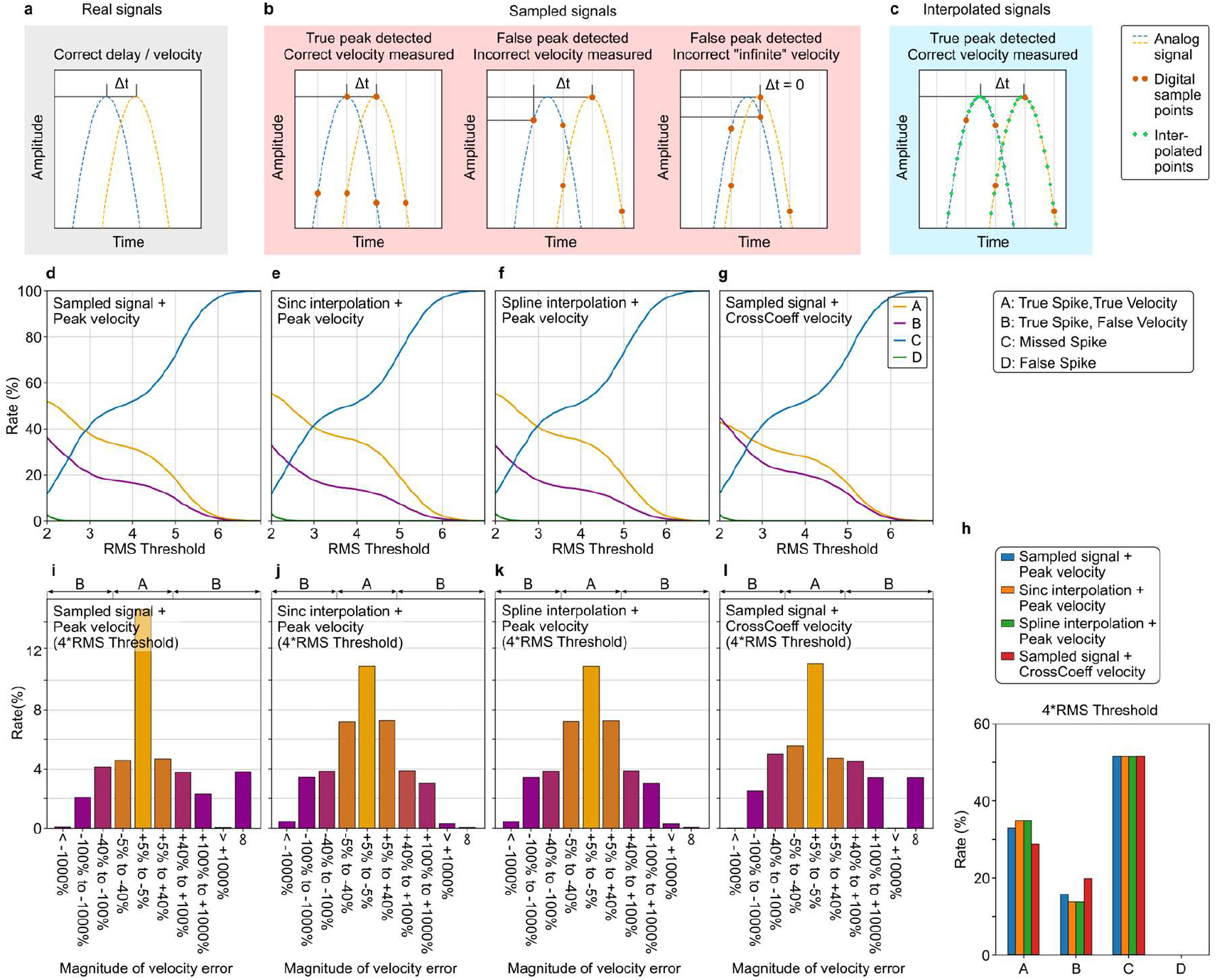
Spike velocity estimation method performance. (a–c) Diagrams summarising peak detection and velocity estimation scenarios. Dashed lines represent analogue signals. Orange circles represent digital sampled signals, which can result in incorrect velocity estimations across the two peaks. Green squares represent upsampled trace through interpolation, which can decrease the impact of these inaccuracies. (a) Real signal with correct delay (Δt) and velocity measurement. (b) Sampled signals, where true peaks (left) or false peaks (centre, right) may be detected due to sampling rate, leading to potentially erroneous velocity estimates. (c) Interpolated signals allowing for more precise detection of peak delay (Δt) for improved velocity estimation. (d–g) Spike detection and velocity error rates for different signal processing methods across RMS thresholds (stepped at 0.05 increments), comparing peak velocity in sampled signals (d), sinc-interpolated signals (e), spline-interpolated signals (f), and cross-correlation coefficient velocity estimation on sampled signals (g). (h) Comparison of rates for each method at a 4*RMS spike detection threshold. (i–l) Histograms representing magnitude of velocity estimation error of spikes for different methods at a 4*RMS threshold, divided into error categories based on deviation from ground truth, with (i) showing errors for sampled signals, (j) for sinc interpolation, (k) for cubic spline interpolation, and (l) for cross-correlation coefficient estimation. Error groups are colour graded from smaller errors (more orange) to larger, more detrimental errors (more purple), the latter of which includes direction flipping (<-100% error, e.g. sensory misinterpreted as motor) and infinite velocity errors (zero time delay). In panels (d-l) A: True Spike, True Velocity (correctly detected spikes with velocity estimates within ±40% of the true value), B: True Spike, False Velocity (correctly detected spikes but with incorrect velocity estimates), C: Missed Spike (true spikes that were not detected), D: False Spike (detected spikes that did not correspond to true spikes). A key for these is also provided in the figure.

**Figure 2.**
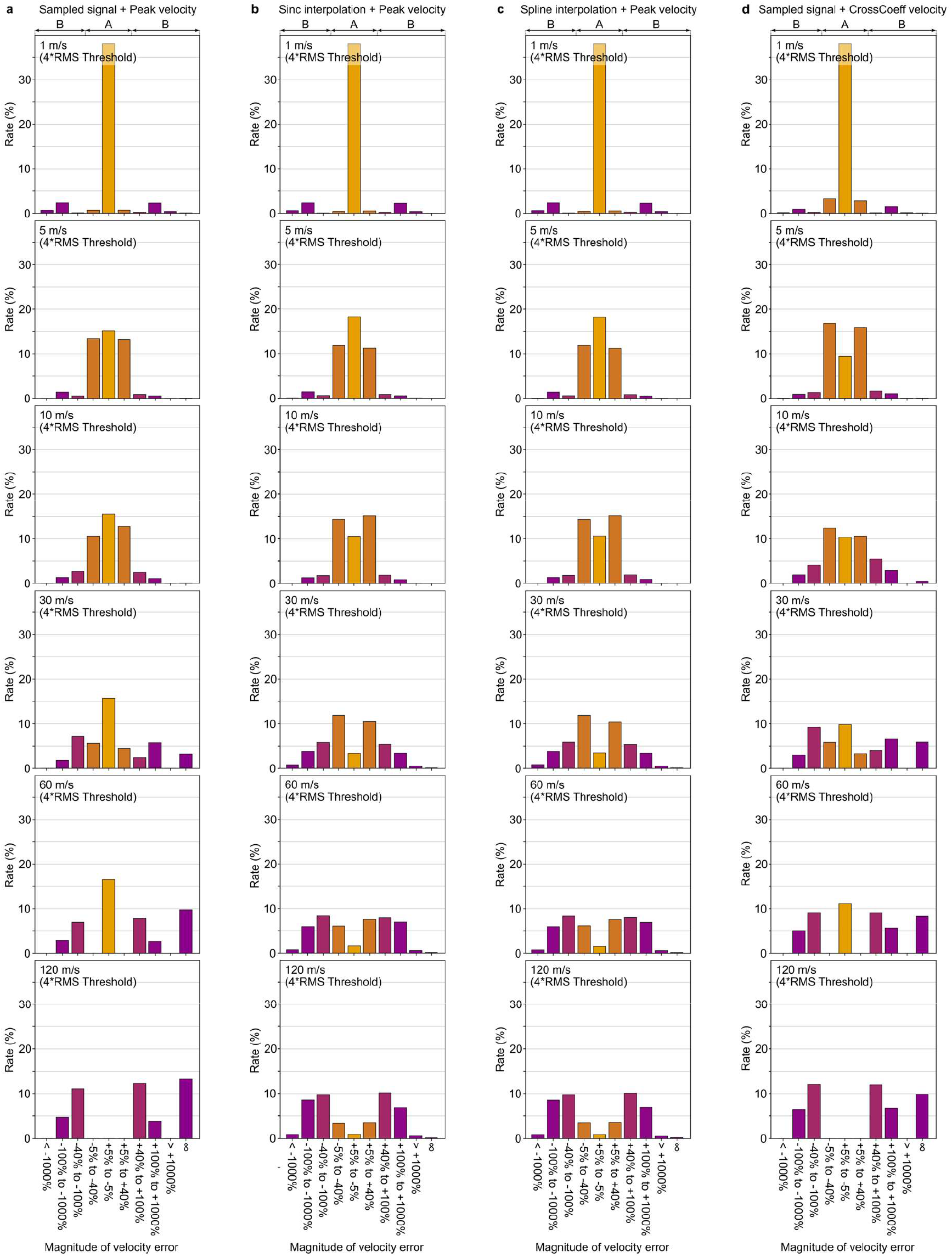
Histograms of magnitude of velocity estimation error of spikes across velocities and methods. (a) Rates of velocity error magnitude for sampled signal with peak velocity estimation, (b) sinc-interpolated signal with peak velocity estimation, (c) cubic spline-interpolated signal with peak velocity estimation, and (d) sampled signal with cross-correlation coefficient velocity estimation. Each row represents a different performance when method is applied to spikes of ground truth velocity: 1, 5, 10, 30, 60, and 120 m/s. Error groups are colour graded from smaller errors (more orange) to larger, more detrimental errors (more purple), the latter of which includes direction flipping (<-100% error, e.g. sensory misinterpreted as motor) and infinite velocity errors (zero time delay).

### 3.4. *In vivo* validation of velocity analysis

Following our results of velocity analysis performance using synthetic data, we sought to test our findings on real nerve recordings. We carried out implantations of parylene-C ultraconformable cuff electrodes into nerves controlling arm function (median, ulnar, and radial nerves) of rats (Fig. 6a) similar to those previously used [12]. We performed recordings of nerve activity while animals were placed and walked along a platform, yielding datasets with both high SNR spike bursts and periods with no visible spikes. Each cuff electrode consisted of rings of microelectrodes at their proximal and distal ends, with an inter-electrode distance of 2 mm, similar to those used in our earlier simulations. This arrangement allowed us to reference the signal of each microelectrode to the average of all of the microelectrodes within the proximal or distal rings, thereby eliminating correlated noise while leaving any proximal-distal time delays in the recorded nerve signal unaltered. This technique, sometimes referred to as common average reference, is often used in other fields to effectively eliminate noise of neural recordings [30].

We first focused on applying the earlier developed cross-correlation velocity analysis to the *in vivo* recordings. We identified a recording containing no visibly apparent spikes (Fig. 6b). While this recording was acquired when the animal was exploring the behavioural chamber and from a somatic nerve of the forelimb (ulnar nerve), no distinct bursts of spike activity were seen by eye within the recording. We evaluated the cross-correlation between proximal and distal recordings and plotted its values over time (Fig. 6b). As earlier, we identified the appropriate averaging window length at which distinct velocity signals could be seen. We found that when using a long (5 s) averaging window we observed the conduction of both afferent and efferent signals of different velocities (seen as horizontal orange bands in the heatmap plot), as well as how these changed over the duration of the recording. In contrast, recordings containing high SNR spike bursts yielded cross-correlation plots with easily identifiable activity at different velocities even with short (200 ms) averaging windows (Fig. 6c).

We examined at which averaging window widths velocity signal patterns could be observed to appear in the previously used low SNR recording. Focusing on an arbitrarily chosen 400 ms fragment of a recording, we found that distinguishable bands of activity in cross-correlation heatmap plots began to be observed with averaging windows of 50 to 100 ms (Fig. 6d). These bands of activity were longer than the averaging windows used to produce the plots, suggesting that these were not an artefact of data averaging over time. Shorter windows of 5 to 10 ms yielded no identifiable bands of activity at any particular delay value. Examining individual cross-correlation plots for a specific timepoint at these averaging window widths highlighted how peaks in cross-correlation only become identifiable above certain window widths – 100 ms in the case of the specific timepoint examined (Fig. 6e). As even longer averaging windows are used, these peaks in cross-correlation became progressively more distinct, while the delays at which they appeared remained generally consistent (Supplementary Fig. 7). Notably, the amplitude of the correlation coefficient peaks also increased with averaging window width, indicating that larger amounts of signal with various time delays were being included, consistent with our earlier simulations (Fig. 3).

**Figure 6.**
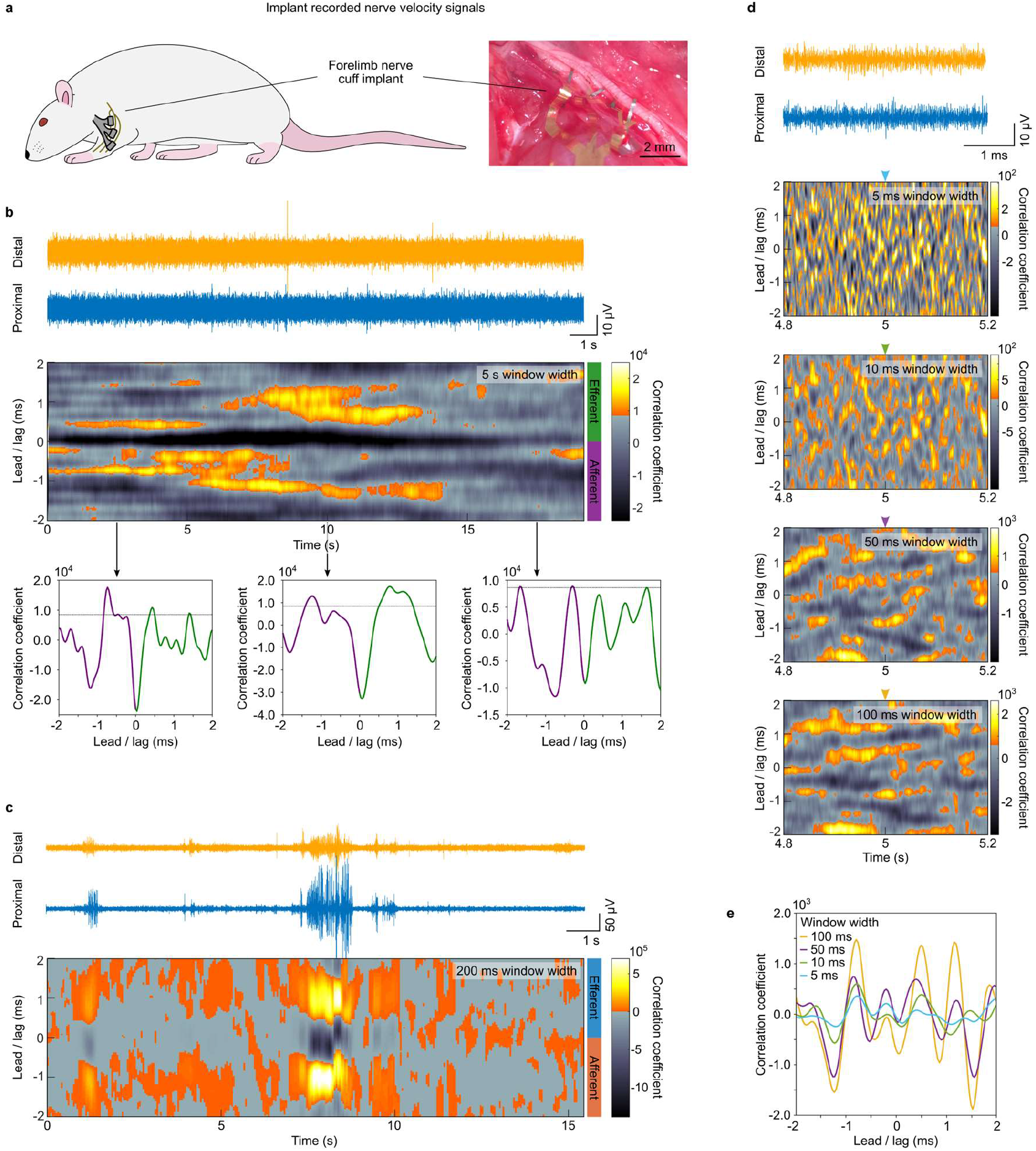
*In vivo* application of cross-correlation velocity analysis. a) Diagram and picture of experimental setup. An ultraconformable parylene-C cuff device is implanted into the median, radial and ulnar nerves innervating the forearm of a rat. b) Recorded traces (top) and cross-correlation heatmap plot (middle) for a proximal and distal electrode in an ulnar nerve cuff for a low SNR activity recording. Cross-correlation heatmap plot is generated with an averaging window of 5 seconds. Lead / lag values correspond to efferent activity (+2 ms to 0 ms) and afferent activity (−2 ms to 0 ms). Plots for cross-correlations at t = 2.5 s, 10 s, and 17.5 s are also provided (bottom). c) Recorded traces and cross-correlation heatmap plot in a median nerve cuff recording high SNR activity. d) Recorded traces and cross-correlation heatmap plots for t = 4.8 to 5.2 s from the same recording used in panel (b), with a range of averaging windows used. Coloured arrowheads indicate cross-correlation values used in panel (e). e) Cross-correlation plots for t = 5 s from heatmap plots in (d). Distinct horizontal bands (d) and peaks (e) can be seen to emerge with longer averaging windows. Transition from grayscale to orange colour scheme in all heatmap plots occurs at the 2*RMS threshold in cross-correlation. This same threshold is indicated by a dashed line in cross-correlation plots in panel (b) (bottom).

Finally, we compared the performance of the various approaches earlier tested for the estimation of spike velocity. In nerve recordings with visible spike activity (Fig. 7a), all four methods yielded similar distributions of spike velocities for the 94-109 spikes co-detected in the +2 to -2 ms delay range (+1 to -1 m/s velocity). The four spike velocity analysis approaches identified similar velocity distributions (Fig. 7b-e), with the biggest difference seen in the ‘Sampled signal + CrossCoeff velocity’. Furthermore, both this group and ‘Sampled signal + Peak velocity’ identified a similar number of spikes with infinite velocity (6 and 5 spikes – 6.4% and 4.6% of all spikes, respectively), while methods including an interpolation step identified none. Furthermore, the two methods including an interpolation step showed identical distributions. Notably, ‘Sampled signal + Peak velocity’ showed a very similar distribution to the interpolation-including methods, with minor differences seen mostly at the fastest spike velocities (between -0.15 and +0.15 ms, equivalent to >13 m/s). These results across methods match the results seen in our earlier simulations (Fig. 4).

**Figure 7.**
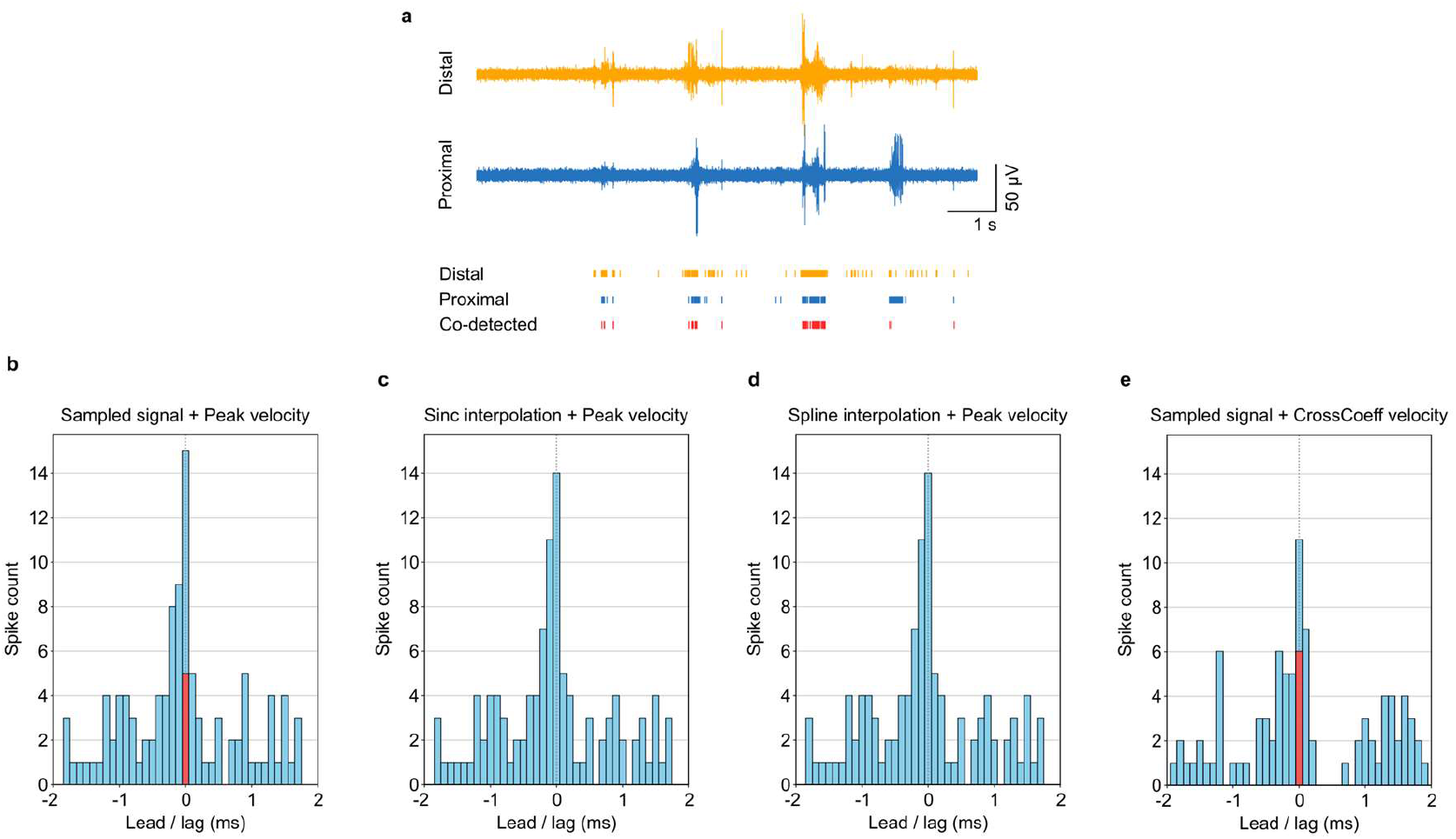
*In vivo* application of spike velocity analysis. a) Recorded traces from proximal and distal cuff electrodes showing spike activity (top). Detected spikes using 4*RMS threshold of trace amplitude for distal and proximal cuff traces, as well as spikes detected simultaneously on both (co-detected) (bottom). b-e) Histogram quantification of spike intervals between co-detected spikes for Sampled signal + Peak velocity (b), Sinc interpolation + Peak velocity (c), Spline interpolation + Peak velocity (d), and Sampled signal + CrossCoeff velocity (e). Histogram bins have a width of 0.1 ms. Within the smallest lead/lag bin (−0.05 to +0.05 ms), spikes with exactly 0 ms lead/lag are highlighted in red. These correspond to spikes with a velocity calculated as infinite.

## 4. Discussion

In this study, we have developed novel tools for analysing signal and spike velocity in nerve cuff recordings, evaluating their performance *in silico* and validating their function *in vivo*. A key limitation of all signal processing is SNR. In the context of neural interfaces recording neural spikes, spikes with an SNR below ∼3 are obscured by the baseline noise, making them inaccessible to detection and processing. We find that in nerve cuff recordings, cross-correlation analysis provides a way to detect these low-amplitude spikes. Specifically, by increasing the window length for cross-correlation, we amplify the cumulative additive effects of spikes, thereby enhancing the effective signal-to-noise ratio. This approach allows for the recovery of neural signals that are not detectable in shorter recordings due to the masking effect of noise. Through synthetic nerve signal simulations, we show that spikes with amplitudes below 1x RMS of the noise with recordings in the tens of seconds. Moreover, our *in vivo* experiments validate this finding, demonstrating that longer averaging windows in cross-correlation analysis can reveal underlying neural activity in peripheral nerve recordings that appear to lack visible spikes. While such long averaging windows may not be suitable for neural interface applications requiring low latencies, many nerve-dependent processes are expected to be driven by activity over longer periods of time. The ability to detect these low-amplitude signals has profound implications for both neuroscience research and clinical neurotechnology applications. In research settings, this enhanced detection capability enables a more comprehensive mapping of neural activity, including signals from small-diameter fibres that are crucial for functions like chronic pain but are traditionally difficult to record due to their low amplitudes. Clinically, improved detection of subtle neural signals can facilitate earlier diagnosis and monitoring of peripheral neuropathies and neurodegenerative diseases, where small changes in nerve activity may precede clinical symptoms. Additionally, this method can enhance the performance of neuroprosthetic devices and closed-loop neuromodulation therapies by providing more accurate and detailed feedback from the nervous system, particularly those relying on slow processes such as bladder state or immune system modulation, leading to more precise interventions and better patient outcomes.

When instead focusing on higher amplitude spikes, our comparative analysis of different spike velocity estimation methods performance was overall good across methods but depended strongly on spike velocity. We found that the simplest method: peak velocity estimation applied directly on the sampled signal is effective for lower velocities but is inherently limited by the digital sampling rate and electrode spacing. To address this, we employed interpolation methods, which increased the detectable velocity range beyond the inherent limitations of the sampling rate, but introduced increased susceptibility to noise leading to increases in misclassifications for certain velocities. Overall, the best choice between these methods ultimately appears to depend on the specific goals of the researcher or clinician. If detecting high-velocity nerve signals is crucial—for example, in studies focusing on fast-conducting motor neurons or large-diameter sensory fibres involved in proprioception—then interpolation will likely be an optimal choice. Conversely, if the priority is obtaining highly accurate velocity estimates and precise directionality for slower fibres, or the latency or computational burden introduced by interpolation is incompatible with the application, then relying on the sampled signal without interpolation may be preferable.

While our specific approach to nerve signal simulation offers valuable insights, it is important to recognize the limitations that come with using a simplified synthetic nerve signal model. We intentionally designed a model with simplified spike shapes, discrete velocities, and no modelling of the effect of axon-electrode distance or axon type on spike amplitude to facilitate a clearer understanding of the tested signal processing approaches. This simplification, however, means that certain aspects of biological nerve signals were not fully represented in our simulations. In reality, spike shapes are not always consistent and may vary across different fibres, with some spikes even overlapping in time, adding complexity to spike detection and analysis. Furthermore, the amplitude of recorded signals depends both on the axon properties as well as on the distance between the axon and the recording electrode. Our study focused on the accuracy and capabilities of velocity analysis methods in nerves. However, factors such as axon heterogeneity and distribution, and how this may be reflected in the analysis, may be of interest to explore in future work to further understand and refine these tools.

Another limitation important to discuss in our simulation approach is noise. In our simulations, we assumed that noise was entirely random and uncorrelated across electrodes, generated by adding separate filtered white noise to either electrode signal. However, in *in vivo* recordings, noise sources are often more complex and will likely not be entirely uncorrelated across electrodes. For example, biological noise can arise from various physiological processes, such as muscle activity and other neural signals, which can introduce correlated noise components across recording sites. Additionally, environmental noise, electrolyte-electrode noise, and thermal noise can contribute to the overall noise spectrum [31]. In practice, noise correlated across electrodes can be eliminated using techniques such as common average referencing [30], as we have employed in our *in vivo* recordings. While we cannot determine whether this eliminates the entirety of correlated noise in electrode recordings, this technique strongly suppresses these while leaving uncorrelated components such as spikes unaffected, indicating that our *in silico* findings should apply to *in vivo* recordings.

Overall, our study shows that both cross-correlation and spike velocity analysis represent powerful approaches to extract velocity information from nerve signals recorded with cuff electrodes. While methods to determine nerve signal velocities already exist [32–37], these methods are typically either applied over larger amplitude evoked compound action potentials and require long arrays of longitudinally placed nerve electrodes. The methods here presented can be implemented on compact nerve cuff designs, requiring minimal data processing, and providing robust results even in challenging environments with dense spontaneous nerve activity such as freely-moving awake recordings. Furthermore, the presented correlation-coefficient method represents to our knowledge the first method to detect nerve activity through signal velocity even in very low SNR environments tested *in vivo*. The advances presented in our work represent powerful tools for nerve recording analysis with applications in both research and clinical translation.

## 5. Conclusion

In this study, we have introduced innovative methods for analysing signal and spike velocity in nerve cuff recordings, demonstrating their effectiveness both *in silico* and *in vivo*. By employing cross-correlation analysis with extended window lengths, we successfully detected and characterised the velocities of low-amplitude neural signals that are typically obscured by noise. In signals containing higher amplitude spikes, we find that spike interval analysis provides a useful tool to characterise spike velocity, applied with or without interpolation depending on for which velocities accuracy should be prioritised. Our findings highlight the potential of these methods to enhance the detection and analysis of nerve signals, representing a valuable advance for neuroscience research and clinical neurotechnology applications.

## Data availability

The data generated in this study and supporting its results have been deposited in the Figshare database under accession code https://doi.org/10.6084/m9.figshare.27687309 [38].

## Code availability

MATLAB and Python code used in data processing and analysis is accessible in the following repository: https://doi.org/10.6084/m9.figshare.27694590.

## Acknowledgements

A.C.L. acknowledges support from the University of Cambridge for a Borysiewicz Interdisciplinary Fellowship and the Wellcome Trust for a Junior Interdisciplinary Fellowship. A.J.B. acknowledges support from his Cross-disciplinary Fellowship (LT000034/2020-C) from the Human Frontier Science Program (HFSP) Organization. R.R.S. acknowledges support from the EPSRC (EP/S022139/1). This work was funded by the ECH2020 FUTURE & EMERGING TECHNOLOGIES (FET) projects BrainCom (732032) and MITICS (964677), and by CloseNIT, an EPSRC/MRC funded Network+, grant code EP/W035057/1.

## Ethical statement

All in vivo procedures were done in accordance with the UK Animals (Scientific Procedures) Act, 1986. Work was approved by the Animal Welfare and Ethical Review Body of the University of Cambridge and was approved by the UK Home Office (project license number PP5478947).

## Author contributions

J.K. carried out *in silico* work. A.C.L. carried out *in vivo* work. A.J.B. carried out the fabrication of devices. J.K., R.R.S. and A.C.L. developed the concept for the analysis techniques. G.G.M. and A.C.L. supervised the work. J.K. and A.C.L. wrote the manuscript. All authors contributed to the review and revision of the manuscript.

## Competing interests

The authors declare no competing interests.

## Notes

### Competing Interest Statement

The authors have declared no competing interest.

