## Supplementary Information for "Development of novel signal and spike velocity analysis tools in peripheral nerve cuffs"

a)

$$\text{spike\_signal}(t) = 10 * \left(-1.8e^{-3(t-3.5)^2} + 0.3e^{-0.4(t-5.3)^2} + 0.1e^{-0.4(t-5)^2}\right)$$

b)

$$\text{spike\_signal} = \frac{\text{spike\_signal}}{\sqrt{\text{mean}(\text{spike\_signal}^2)}} * \text{spike\_rms}$$

**Supplementary Figure 1: Spike Signal Modelling Function**. a) Mathematical formulation used to model extracellularly recorded spike shapes in our analysis. The spike signal, denoted as spike_signal(t), was generated using the equation (a) to approximate extracellular spike shapes. b) To ensure the signal matched a target root-mean-square (RMS) value, a normalization and scaling step was applied as per the indicated equation. Each exponential term in the formula modelled a specific component of the spike shape, allowing for a realistic approximation of extracellularly recorded spikes. The scaling step adjusted the generated signal to achieve the desired RMS value, ensuring consistency in signal amplitude across different simulations.

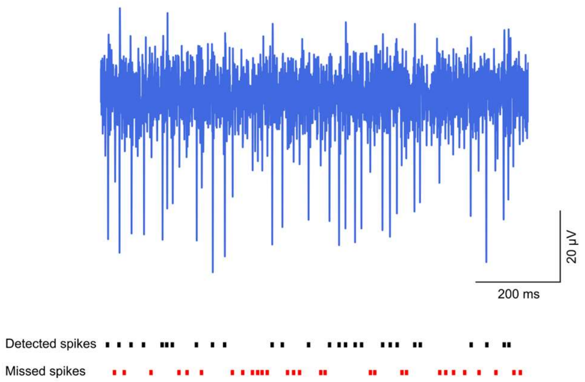


**Supplementary Figure 2. Example of single synthetic signal trace with ground truth spikes and detection outcomes used in spike velocity estimation evaluation.** (Top) 1-second simulated nerve signal trace containing approximately 60 ground truth spikes. (Bottom) ground truth spikes. Black bars represent detected spikes for which a corresponding peak was found in the second signal, allowing for velocity estimation. Red bars indicate missed spikes, which were either not identified as peaks in the first signal based on a 4*RMS threshold or lacked a corresponding peak in the second signal.

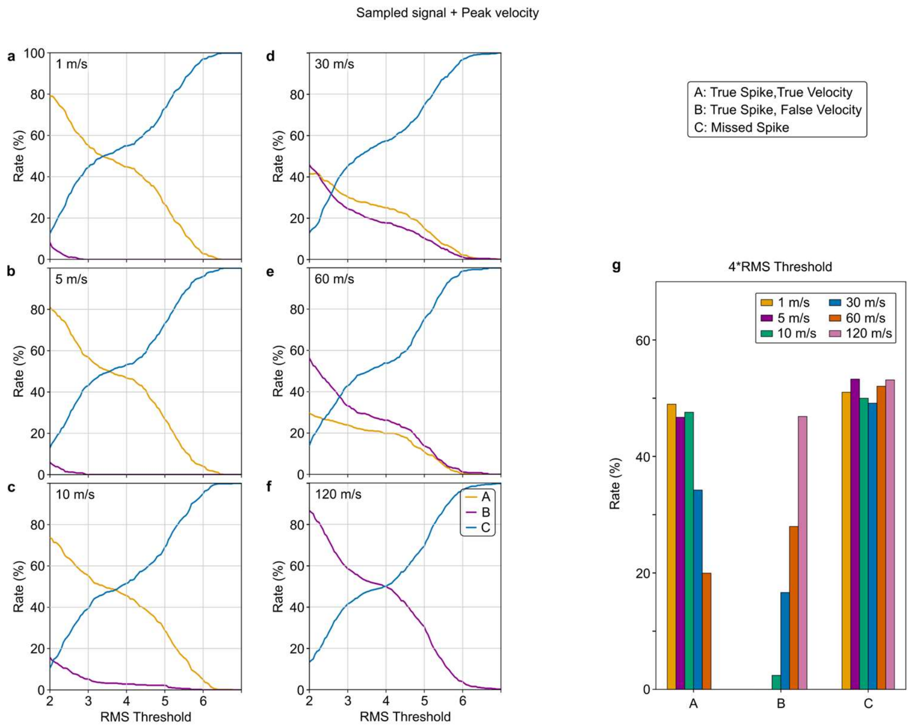


**Supplementary Figure 3**. **Influence of signal threshold on spike detection performance at different velocities using peak velocity estimation.** Panels (a-f) illustrate the rates of correctly detected spikes with accurate velocity estimation (A - True Spike, True Velocity), correctly detected spikes with incorrect velocity estimation (B - True Spike, False Velocity), and missed spikes (C - Missed Spike) across a range of RMS thresholds (stepped at 0.05 increments) for various spike velocities: (a) 1 m/s, (b) 5 m/s, (c) 10 m/s, (d) 30 m/s, (e) 60 m/s, and (f) 120 m/s. At lower velocities (1, 5, and 10 m/s), accurate detection remains relatively stable as RMS thresholds increase. However, for higher velocities (30 and 60 m/s), there is a marked decline in detection accuracy as RMS thresholds rise, with spikes at 120 m/s being completely undetectable due to limitations in sampling rate and signal resolution. Panel (g) summarizes detection outcomes for each velocity at a 4*RMS threshold, highlighting that while low-velocity spikes (1 and 5 m/s) maintain high detection accuracy, higher velocities are increasingly prone to misclassification or missed detections, culminating in the inability to detect 120 m/s spikes altogether. Category D - False Spikes is not depicted here, as false spikes cannot be attributed to a specific ground truth velocity due to their lack of association with an actual ground truth spike. While these false spikes can be shown in the aggregated Figure 4d, they cannot be represented in velocity-specific panels.

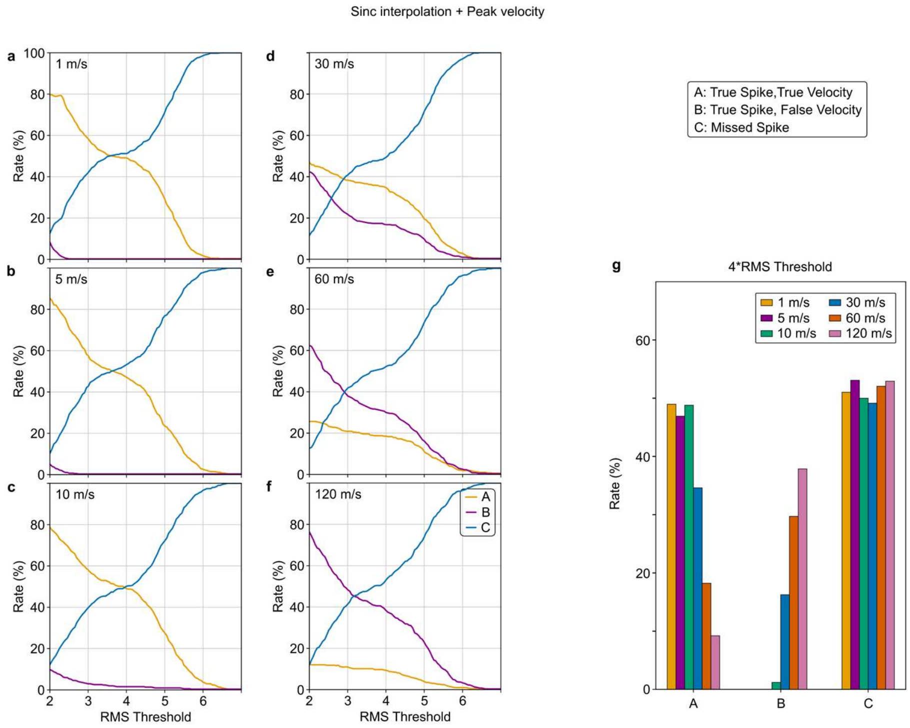


**Supplementary Figure 4. Influence of signal threshold on spike detection performance at different velocities using sinc interpolation with peak velocity estimation.** Panels (a-f) display detection rates for true spikes with correct velocity estimation (A - True Spike, True Velocity), true spikes with incorrect velocity estimation (B - True Spike, False Velocity), and missed spikes (C - Missed Spike) across a range of RMS thresholds (stepped at 0.05 increments) for different velocities: (a) 1 m/s, (b) 5 m/s, (c) 10 m/s, (d) 30 m/s, (e) 60 m/s, and (f) 120 m/s. Sinc interpolation enhances temporal resolution by filling gaps between discrete sampling points, allowing accurate detection of spikes at higher velocities, which were previously undetectable on the non-interpolated signal due to sampling limitations. For lower velocities (1, 5, and 10 m/s), sinc interpolation performs similarly to the non-interpolated approach, maintaining high detection accuracy across a range of thresholds. However, at 30 m/s and 60 m/s, sinc interpolation prevents infinite velocity detections that occurred in the non-interpolated model, enabling more reliable detection. At 120 m/s, sinc interpolation allows some detection, although misclassification and missed detections remain significant due to residual noise effects. Panel (g) summarizes detection outcomes at a 4*RMS threshold across all velocities, highlighting that sinc interpolation improves detection rates for higher velocities relative to the non-interpolated approach. Category D - False Spikes is not depicted here, as false spikes cannot be attributed to a specific ground truth velocity due to their lack of association with an actual ground truth spike. While these false spikes can be shown in the aggregated Figure 4e, they cannot be represented in velocity-specific panels.

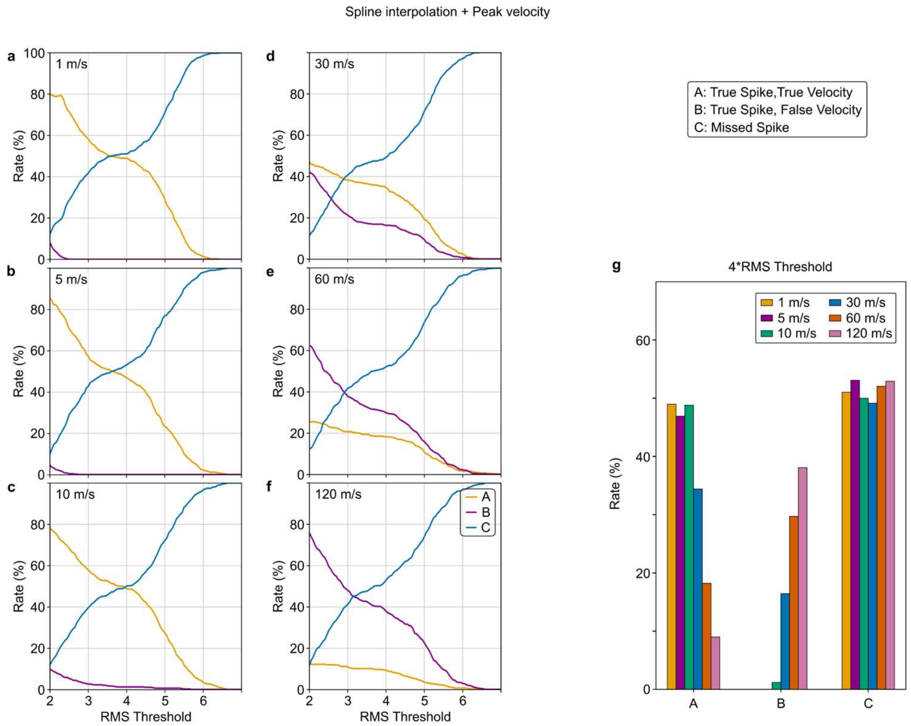


**Supplementary Figure 5. Influence of signal threshold on spike detection performance at different velocities using cubic spline interpolation with peak velocity estimation.** Panels (a-f) display detection rates for true spikes with correct velocity estimation (A - True Spike, True Velocity), true spikes with incorrect velocity estimation (B - True Spike, False Velocity), and missed spikes (C - Missed Spike) across a range of RMS thresholds (stepped at 0.05 increments) for different velocities: (a) 1 m/s, (b) 5 m/s, (c) 10 m/s, (d) 30 m/s, (e) 60 m/s, and (f) 120 m/s. Similar to sinc interpolation, cubic spline interpolation enhances temporal resolution by generating a continuous signal from discrete sampling points, enabling the detection of spikes at higher velocities that are typically undetectable in non-interpolated signals due to sampling rate limitations. For lower velocities (1, 5, and 10 m/s), cubic spline interpolation maintains a high rate of correct detection across various thresholds, showing similar performance to sinc interpolation. At 30 m/s and 60 m/s, cubic spline interpolation prevents infinite velocity detections that are observed in non-interpolated models, resulting in more reliable detection. However, at 120 m/s, cubic spline interpolation allows for partial detection, though misclassification and missed detections remain prominent due to residual noise influences. Panel (g) provides a summary of detection outcomes at a 4*RMS threshold for each velocity, demonstrating that cubic spline interpolation enhances detection rates for higher velocities compared to the non-interpolated signal. Overall, cubic spline interpolation performs almost identically to sinc interpolation, offering similar improvements in high-velocity detection and preventing infinite velocity errors while maintaining accuracy at lower velocities. Category D - False Spikes is not depicted here, as false spikes cannot be attributed to a specific ground truth velocity due to their lack of association with an actual ground truth spike. While these false spikes can be shown in the aggregated Figure 4e, they cannot be represented in velocity-specific panels.

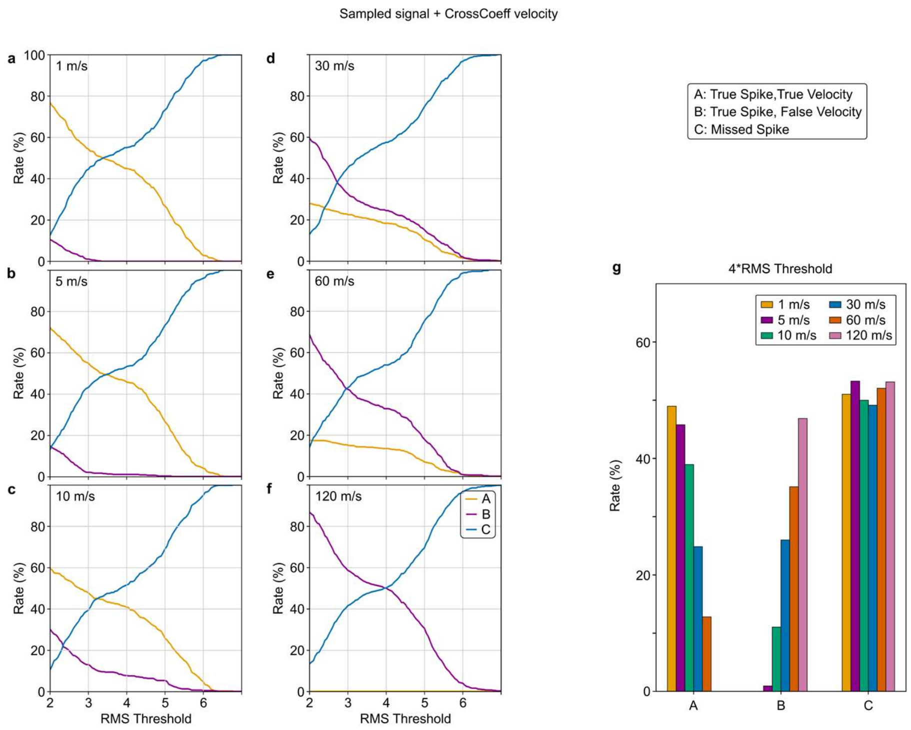


**Supplementary Figure 6. Influence of signal threshold on spike detection performance at different velocities using cross-correlation coefficient velocity estimation on the sampled signal.** Panels (a-f) display detection rates for true spikes with correct velocity estimation (A - True Spike, True Velocity), true spikes with incorrect velocity estimation (B - True Spike, False Velocity), and missed spikes (C - Missed Spike) across a range of RMS thresholds (stepped at 0.05 increments) for different velocities: (a) 1 m/s, (b) 5 m/s, (c) 10 m/s, (d) 30 m/s, (e) 60 m/s, and (f) 120 m/s. Unlike the peak velocity method, cross-correlation coefficient estimation exhibits greater sensitivity to noise, leading to a higher rate of incorrect detections at velocities beyond 10 m/s. At lower velocities (1 and 5 m/s), the method achieves relatively high detection accuracy across various thresholds. However, performance declines significantly for higher velocities. Specifically, for velocities of 30 m/s and above, cross-correlation coefficient estimation results in a notable increase in misclassifications and missed spikes, with 120 m/s showing a complete failure to reliably detect true spikes, attributed to the limitations of sampling rate similar to the peak velocity method used on the sampled signal. Panel (g) provides a summary of detection outcomes at a 4*RMS threshold for each velocity, indicating that while cross-correlation coefficient estimation maintains moderate accuracy at low velocities, it struggles with increased error rates and undetectable spikes at high velocities. Category D - False Spikes is not depicted here, as false spikes cannot be attributed to a specific ground truth velocity due to their lack of association with an actual ground truth spike. While these false spikes can be shown in the aggregated Figure 4e, they cannot be represented in velocity-specific panels.

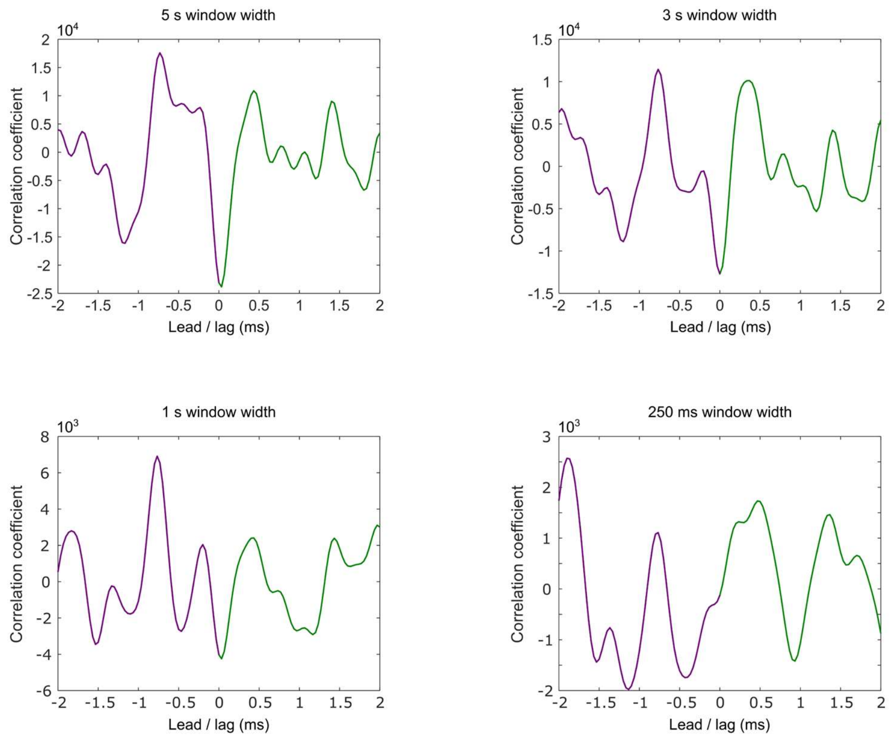


**Supplementary Figure 7. Effect of averaging window width on velocity pattern detectability in cross-correlation analysis of *in vivo* peripheral nerve recordings.** Cross-correlation plots for varying averaging windows demonstrate how peak distinctness and correlation coefficient amplitude increase as the window lengthens, while the delays remain generally consistent across conditions. Panels show cross-correlation results with different averaging windows: 5 s (upper left), 3 s (upper right), 1 s (lower left), and 250 ms (lower right). Longer averaging windows incorporate more signal with various time delays, leading to clearer and more distinct peaks in cross-correlation, in alignment with trends observed in our synthetic data simulations (Fig. 3).
